# Mature black alder shows intermittent events of high stem methane emissions

**DOI:** 10.1101/2020.09.01.277350

**Authors:** Daniel Köhn, Anke Günther, Ines Schwabe, Gerald Jurasinski

## Abstract

- Tree stems can be a source of the greenhouse gas methane (CH_4_). However, assessments of the global importance are complicated by a lack of research and a high variability between ecosystems. Here, we determined the contribution of emissions from tree stems of mature black alder (*Alnus glutinosa* (L.) Gaertn.) to overall CH_4_ exchange in two temperate peatlands.
- We measured emissions from stems and soils using closed chambers in a drained and a wet alder forest over two years. Further, we studied the importance of alder leaves as substrate for methanogenesis in an incubation experiment.
- Stem CH_4_ emissions were shortlived and occurred only during times of inundation at the wet site. The drained site did not show stem emissions and the soil acted as a small CH_4_ sink. The contribution of stem emissions to the overall CH_4_ budget was below 0.3% in both sites.
- Our results show that also mature black alders intermittently can be a source of CH_4_. However, the low share of stem-mediated CH_4_ emissions in both forests may indicate that this pathway is only of minor relative importance in temperate peatlands.

## Introduction

Tree stems are increasingly recognized as potential surfaces for the exchange of greenhouse gases (GHG) such as methane (CH_4_) (Pangala *et al*., 2013, 2015). CH_4_ is a potent GHG whose globally relevant emission sources are not yet fully understood (Melton et al., 2013; Saunois *et al*., 2016). It is possible that tree stems are a source that has been previously overlooked in global CH_4_ inventories (Pangala *et al*., 2017). Accordingly, emissions from trees have been discussed as a “new frontier in the global carbon cycle” (Barba *et al*., 2018).

Regionally, tree stem emissions can be the most important CH_4_ source (e.g. in the amazon basin, Pangala et al. 2017). However, the magnitude of CH_4_ fluxes from tree stems seems to vary considerably on a global scale with fluxes tending to be lower in temperate (Gauci *et al*., 2010; Pitz *et al*., 2018) than in tropical regions(Pangala *et al*., 2017). The ecosystem share of stem CH_4_ emissions has only rarely been addressed, with the few available studies reporting shares between 27% in a temperate and 87% in a tropical wetland (Pangala *et al*., 2013, 2015).

Although CH_4_ emissions from tree stems could be an emission pathway of global relevance, little information is available on the influence of ecosystem type or of abiotic factors. e.g. water level and soil temperature. Also, the variability of stem emissions between and within individual trees, e.g. caused by stem height, tree age or tree species, has rarely been studied (Barba *et al*., 2018).

Abiotic and biotic factors can influence both CH_4_ production itself and its emission rate from tree stems. Generally, soil temperature, soil water content or water level and physiological activity of the trees are assumed to be the most important influencing factors (Terazawa *et al*., 2015; Barba *et al*., 2019; Schindler *et al*., 2020). Accordingly, a considerable increase in stem CH_4_ emissions may be associated with increasing water tables (Pitz *et al*., 2018; Schindler *et al*., 2020) and increasing soil and air temperatures (Barba *et al*., 2019). This is further backed by mesocosm experiments (Rusch & Rennenberg, 1998; Pangala *et al*., 2014).

The origin of the emitted CH_4_ likely can be either the tree trunk itself or the soil. Several studies found stems of upland trees to be sources of CH_4_ despite standing on very dry soils (Machacova *et al*., 2016; Wang *et al*., 2016; Maier *et al*., 2017; Pitz & Megonigal, 2017). Thus, heartwood rot can be a good indicator for stem CH_4_ emissions (Covey *et al*., 2012; Wang *et al*., 2016). In addition, methanogenesis may even take place in aerobic parts of plant tissue (Bartlett *et al*., 2006; Bižić *et al*., 2020). On the other hand, CH_4_ produced in the soil can enter the tree through the roots and may be transported through the sap or through air-filled spaces in the trunk (Schröder, 1989). This is supported by several studies finding higher CH_4_ emissions in the lowest parts of the trunk (Terazawa *et al*., 2007; Barba *et al*., 2019; Schindler *et al*., 2020).

Trees like black alder (*Alnus glutinosa* (L.) Gaertn.) additionally have the ability to transport oxygen into their rhizosphere and thereby stimulate methantrophy (Joabsson & Christensen, 2001). In addition, alders predominantly grow on peat soils in temperate regions, which themselves have a high potential for CH_4_ emissions (Turetsky *et al*., 2014).

Black alder forests roughly cover 5% of all central European forests (Claessens *et al*., 2010). In north-eastern Germany black alder is the second most important deciduous tree species after the European beech (*Fagus sylvatica* L.). In addition, the importance of black alder could increase further since it has a high potential for sustainable silviculture on rewetted peatlands (Schäfer *et al*., 2005).

Here, we study patterns of CH_4_ emissions through stems of mature black alders and relate them to soil CH_4_ emissions to evaluate the share of CH_4_ passing through trees in two temperate alder forests over two years. One of the studied forests is currently drained while the other is undrained (i.e. wet). Further, we studied the possible effects of leaf fall on soil CH_4_ emissions by determining CH_4_ production potentials in an incubation experiment. Our hypotheses are:

1. CH_4_ emissions from stems of black alder contribute significantly to the overall CH_4_ budget.
2. Absolute values of tree stem CH_4_ emissions of the wet site are larger than at the drained site, while relative contributions are higher at the drained site due to low or no emissions from the soil.
3. Events like inundation, leaf fall or leaf out significantly increase stem CH_4_ emissions, because inundation increases CH_4_ production, leaf fall provides substrate for methanogenesis and leaf out increases rates of internal transport in the tree due to higher transpiration.

## Material and Methods

### Study sites

The drained and wet black alder forest sites are located in north-eastern Germany (54° 7’ 36.27” N, 12° 28’ 55.5” E) sharing a distance of approx. 1 km. The drained site has been logged for over 200 and most likely drained even earlier. After decades of drainage the wet site has gradually been rewetted since the 1990s with the final water levels establishing in 2004. Both sites are located approx. 40 m.a.s.l. and receive 627 mm of rain on average (DWD, reference period: 1981-2010) with annual mean temperature of 8.5 °C. Both sites are situated in local depressions. The research sites are fenced areas of approx. 12 x 30 m with a boardwalk running along the long side through the center of the short side. Boardwalks were constructed between April and June 2017.

**Abbildung 1 Fig. 1:**
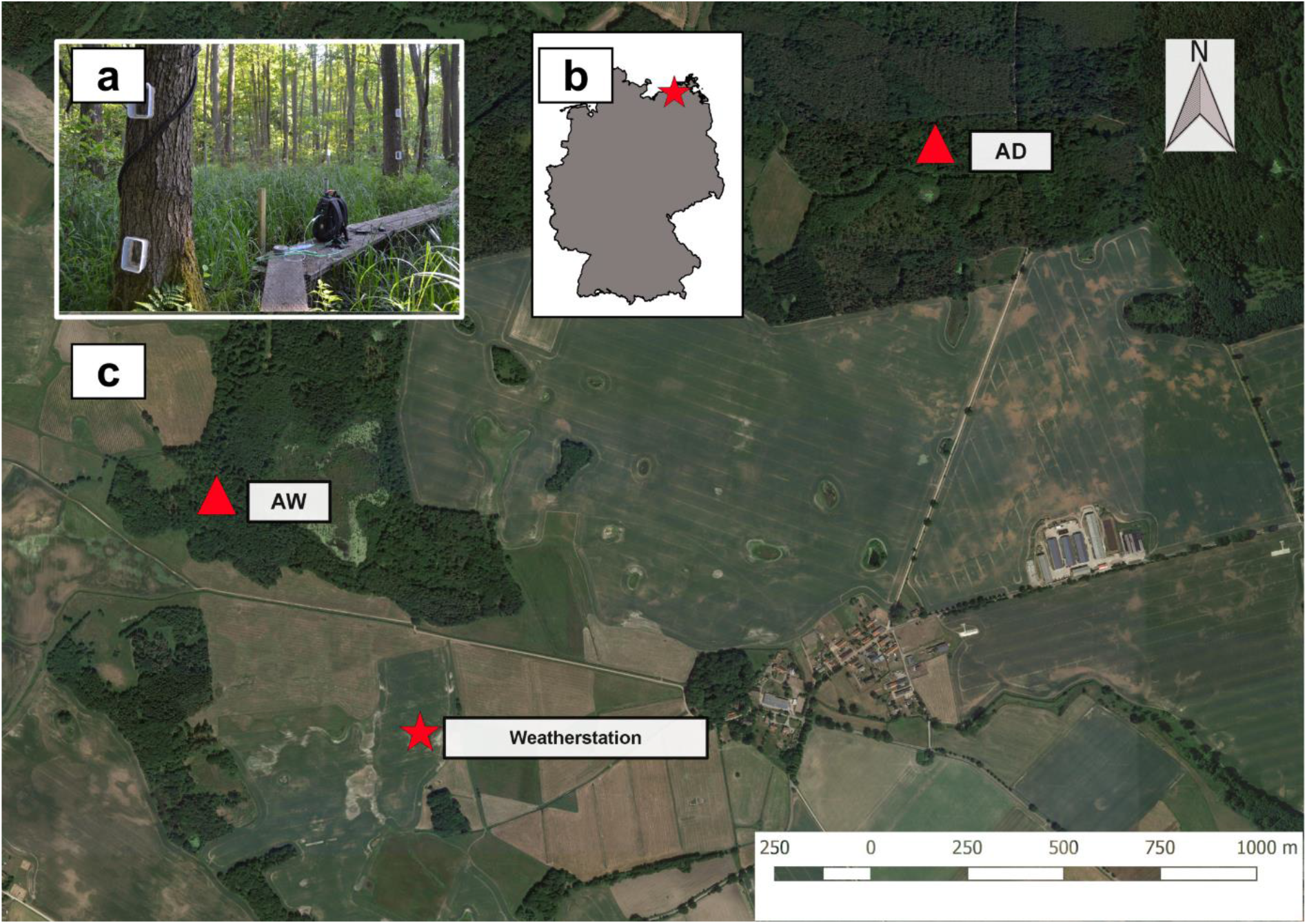
**a)** Setup of tree chambers and Picarro Gas Scouter at AW. **b)** Study location in Germany. **c)** Location of AW, AD and the weatherstation

Black alder (*Alnus glutinosa* (L.) Gaertn.) is the only tree species in the wet site (AW – **A**lder **W**et). The understorey features sedges (*Carex acutiformis* Ehrh.), *C. riparia* Curtis), featherfoil (*Hottonia palustris* L.) and bitter nightshade (*Solanum dulcamara* L.). However, temporal flooding created areas of open water. Peat depth is around 2 m with an average soil pH of around 5.1. Soil carbon and phosporus contents lie at 38 % and 1.4 %, respectively (Jurasinski *et al*., 2020).

The drained site (AD – **A**lder **D**ry) features a mixed stand of black alder and European ash (*Fraxinus excelsior* L.). The understory mainly comprises nettle (*Urtica dioica* L.), elderberry (*Sambucus nigra* L.) and blackberry (*Rubus sp* L.). In spring lesser calendine (*Ranunculus ficaria* (HUDS.)) is dominant on the ground. The peat is very shallow with a depth of approximately 40 cm. Soil pH is 4.5 with soil carbon and phosphorus content being 19 % and 2.7 %, respectively (Jurasinski *et al*., 2020).

### Study setup

We used flow-through non-steady-state chambers to measure gas exchange between the atmosphere and the stem as well as the soils surface, respectively (Livingston & Hutchinson, 1995). We measured soil exchange fluxes in both sites at five different spots with permanently installed PVC collars in the ground (approx. 10 cm depth, radius = 65 cm, height = 0.9 – 1.4 m) (Günther *et al*., 2014). Soil chambers were equipped with three fans at the lid as well as a temperature and humidity sensor. Stem chambers were permanently installed on three trees at three heights above the base of the stem (approx. 0.3 m, 1 m, 2 m) at each site (i.e. 9 chambers per site). We constructed the stem chambers using polypropylene containers from which the bottom was cut out (EMSA^®^ V = 0.001 m^3^). The chambers were mounted on the tree stems with an adhesive synthetic sealant (plastic fermit^®^, Fermit).

### Flux measurements

Measurements were carried out every two weeks over the course of two years between May 3^rd^ 2018 and April 30^th^ 2020. On June 26^th^ 2018 we ran an 18-hour campaign to determine differences in CH_4_ emissions from tree stems during day and night. In this campaign every stem chamber was measured every 45 minutes.

For measurements of soil GHG exchange, the flexible side walls of the chamber were firmly attached to the PVC collar by a rubber band. For stem chamber measurements, the chamber was closed using a lid with a rubber gasket. Inlet and outlet tubes (PVC, inner diameter: 3.4 cm) connected the chambers to a portable gas analyser. CO2 and CH_4_ concentrations inside the chamber were measured by laser spectrometers (Ultra-Portable Greenhouse Gas Analyzer, Los Gatos Research ©, and GasScouter, Picarro ©) with a measurement frequency of 1 Hz. Enclosure time for soil and stem chambers was 5 and 3 minutes, respectively. A shorter enclosure time for the stem chambers was chosen due to the much smaller headspace volume (0.002 m^3^ vs. 0.31 m^3^). Flux estimation was performed using R 3.6.4 (R development core team, 2020). For flux estimation we used an updated version of the *fluxx*() function of package *flux* for R (Jurasinski *et al*., 2014). The slope was calculated between each concentration measurement (one per second). Subsequently the median of the slope values was taken as an estimate for the gas flux (i.e. median based regression (Siegel, 1982)).

### Additional measurements

A weather station 500 m SW of the wet site (CR300, Campbell Scientific^®^) recorded air temperature, wind velocity, precipitation and radiation (PPFD). At both sites Onset^®^ HOBO Pendant soil temperature loggers were used at three different locations in 5 and 15 cm depth with a measurement interval of 15 min. Water levels were logged continuously with a SEBA^®^ CS457 dipper at 1 min intervals.

### Methane budgets

Since we were not able to model methane fluxes using ancillary data, budgeting had to be based on a statistical approach adapted from Günther *et al*., (2017).We used the area-under-curve function (auc.mc) from the R package *flux* (Jurasinski et al. 2014) to integrate flux values over time by linear interpolation. For each measurement day, one flux value per flux subset (stem or soil) and each site (drained or wet) was randomly chosen. This was repeated 100 times to obtain 100 different flux time series. Then, the area-under-curve was calculated 100 times for each flux time series, each time leaving out one flux value (jackknife method), leading to a total of 10 000 different CH_4_ balances. For the final CH_4_ balances per site and flux subset we calculated the average and standard deviation of all balances.

For determining the share of stem and soil mediated CH_4_ emissions, we projected the stem emissions of the lower 2.2 m of the stem onto the base area of the respective tree trunks. The upper limit of 2.2 m was chosen because we assumed that no CH_4_ was emitted from the tree trunks above that height (see results). First, we measured the stem diameters of all sampled trees once at the stem base and once at 2.2 m height. Then, we calculated the surface area of the lower 2.2 m of each tree (A_Surface_) by using equation (1) that assumes the tree trunk to be a circular conical fustrum:

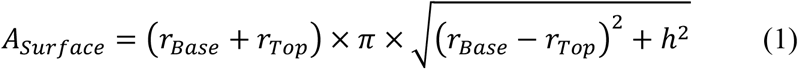

where *r_Base_* and *r_Top_* are the base and top radii and *h* is the height (2.2.m). The base area of each tree trunk was calculated by equation (2):

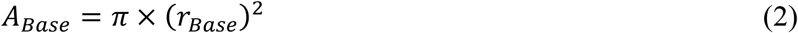

Finally, we calculated the stem CH_4_ flux projected to ground surface (F_Base_) using equation (3)

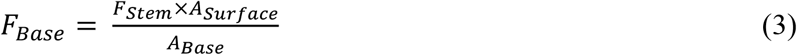

where F_stem_ is the flux estimated from stem chambers relative to the stem area. Because no consensus exists on how to upscale stem fluxes, especially with regards to whether fluxes are upscaled on a stem or soil area basis we aimed at comparing three different approaches:

A1. We used FBase to calculate annual balances and upscaled it with the approx. total base area of all trees per hectare.
A2. We used the daily average stem flux of each tree and multiplied it with the respective surface area of the stem which yields an average flux per tree and day. With this we also calculated annual balances.
A3. We used the overall average of stem fluxes and multiplied it with the total approx. stem surface area of all tree stems and projected it to the period of one entire year.

For each approach we calculated the share of stem emissions with regards to the total emissions of the ecosystem. For A1 and A2 we used the calculated balance for soil CH_4_ emissions for comparison. For A3 the overall average soil CH_4_ flux was taken for comparison. Soil CH_4_ fluxes and soil CH_4_ balances were spatially projected by multiplying with 1 ha excluding the base area of all trees per ha (AD: 273 trees ha^−1^ with 137 m^2^ base area, AW: 311 trees ha^−1^ with 106 m^2^ base area). Upscaling and the calculation of the share of stem CH_4_ emissions were done separately for AD and AW and each study year.

### Incubation experiment

We complemented our field monitoring data with an incubation experiment to study the possible effects of leaf fall on soil CH_4_ emissions. For the incubation experiment soil samples were taken at both sites from the top soil (0-10cm) in January 2020 and stored at approx. 5°C in a fridge for 9 days. Alder leaves were collected in fall 2018 and dried for 3 days at 60°C and subsequently stored in air tight plastic bags. Leaves were ground to 0.5 mm size. The incubation experiment comprised three different anaerobic incubations. We set up three incubation treatments to determine the CH_4_ production potential of soil and leaves separately and in combination. Hence, each incubation container (140 ml, Weck^®^) received either 10 g soil (*Soil_only_*) or 50 mg leaves (*Leaves_only_*) or a combination of 10 g soil and 50 mg leaves (*Soil +Leaves_combination_*). To establish anaerobic conditions 10 ml of de-ionized water were added to the containers, after which the containers were closed with a gas-tight lid. Then, the headspace was purged for 30 seconds with pure N_2_ gas. This procedure was repeated after each sampling. The anaerobic containers remained closed during the entire experiment.

In total, the experiment ran for 34 days. Gas samples were taken at two-day intervals at the beginning and later at four-to six-day intervals. Gas samples were taken from the containers with a 60 ml syringe. Approximately 25 ml headspace gas was added to evacuated 12 ml Labco^®^ Exetainers. The analysis of the gas samples was carried out with a gas chromatograph (Shimadzu Auto System) within three days after sampling.

The production potential of the control series was calculated as follows:

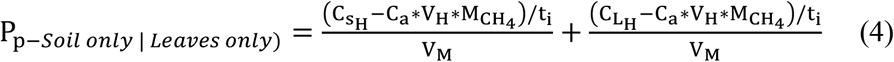

And the production potential of the priming series as follows:

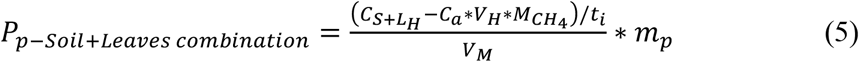

Where C_sH_ and C_LH_ are the CH_4_ concentrations in the headspace samples of the incubated soil and leaves, respectively. C_S+LH_ is the CH_4_ concentration in the headspace of the incubated combination of soil and leaves. C_a_ is the starting concentration of CH_4_ (assumed to be 0), VH the headspace volume, V_M_ the molar volume of CH_4_ (24.7 l mole^−1^ at 25°C), M_CH4_ the molar mass of CH_4_ (16.04 g mole^−1^) and t_i_ the duration of the incubation. The production potential was calculated in relation to the mass of the material that was incubated. Finally, the separate CH_4_ production potentials of *Soil_only_* and *Leaves_only_* taken together were compared with *Soil+Leaves_combination_* in order to prevent weight effects on the CH_4_ production potential.

### Statistical analyses

Statisical analyses and visualizations were carried out with R 3.6.4 (R development core team, 2020). The entire datasets and subsets were tested for normality using Shapiro Wilk tests. We used Kruskal-Wallis tests to test for differences in means of the not normally-distributed data. Relationships between abiotic variables and CH_4_ fluxes were carried out with stepwise multiple linear regressions or exponential linear regressions.

## Results

### Climate

The study period was characterised by a large precipitation deficit together with above average temperatures compared to the respective average values of the climate period 1981–2010 (data from the German Weather Service, DWD). In the first study year (May 2018-April 2019) the total precipitation was 381 mm (60% of long-term average) and 424 mm (67% of long-term average) in the second year (May 2019-April 2020) (Fig. 2). In both years, the deficit was especially prevalent in spring and summer. The average air temperature was 10.4 and 10.9 °C for year one and two, respectively. This is 1.7 and 2.2 °C above the annual mean temperature. The water table was very variable at both the drained and the wet site.

**Abbildung 2 Fig. 2:**
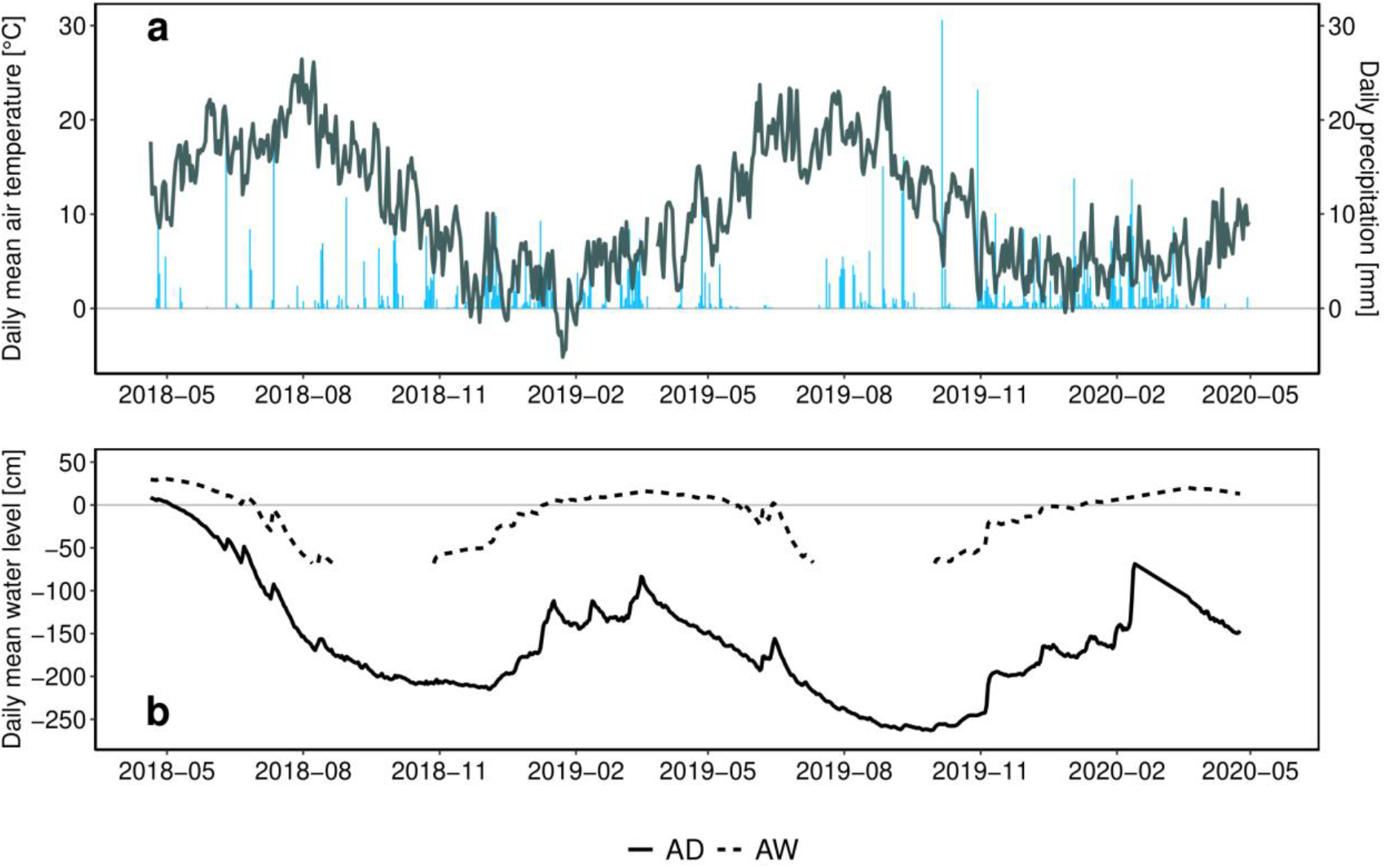
**a)** Daily mean air temperature and daily precipitation. **b)** Daily mean water level in cm below soil surface. Solid line shows AD, dashed line shows AW. Gaps are present because the water level fell below the deepest point of the well.

### Flux measurements

In total, 1167 stem flux and 414 soil flux measurements were conducted. Soil as well as stem CH_4_ fluxes were significantly higher in AW than in AD (p < 2.2*10^−16^ and p = 8.5*10^−4^, respectively). During the study period, AD tended to show CH_4_ uptakeby the soil, with an average flux of −0.07 ± 0.12 mg m^−2^ h^−1^, while CH_4_ exchange was not visible at the tree stems. Meanwhile, CH_4_ emissions were temporarily high both from the soil and the stem surface in AW (average ± SD: 4.8 ± 18.8 mg m^−2^ h^−1^ and 0.07 ± 0.34 mg m^−2^ h^−1^, respectively) without showing a clear seasonal pattern. However, emissions from both stems and soil in AW were very short lived and mostly lasted for only 6-8 weeks a year. The highest stem CH_4_ flux was 4.0 mg m^−2^ h^−1^ (relative to stem area) on May 15^th^ 2018 while the highest recorded soil CH_4_ flux was 132.4 mg m^−2^ h^−1^ on June 29^th^ 2018 (Fig. 3). An intensive measuring campaign over 18 hours of high-frequency measurements showed no diurnal pattern of CH_4_ emissions from the tree stems at AW (data not shown). Overall, CH_4_ emissions from the individual trees sampled in AW differed significantly from each other (p = 3.2*10^−3^). Despite a trend of decreasing stem CH_4_ emissions with increasing height (Fig. 4) the three different height groups were only significantly different at one tree (p = 7.9*10^−3^).

**Fig. 3:**
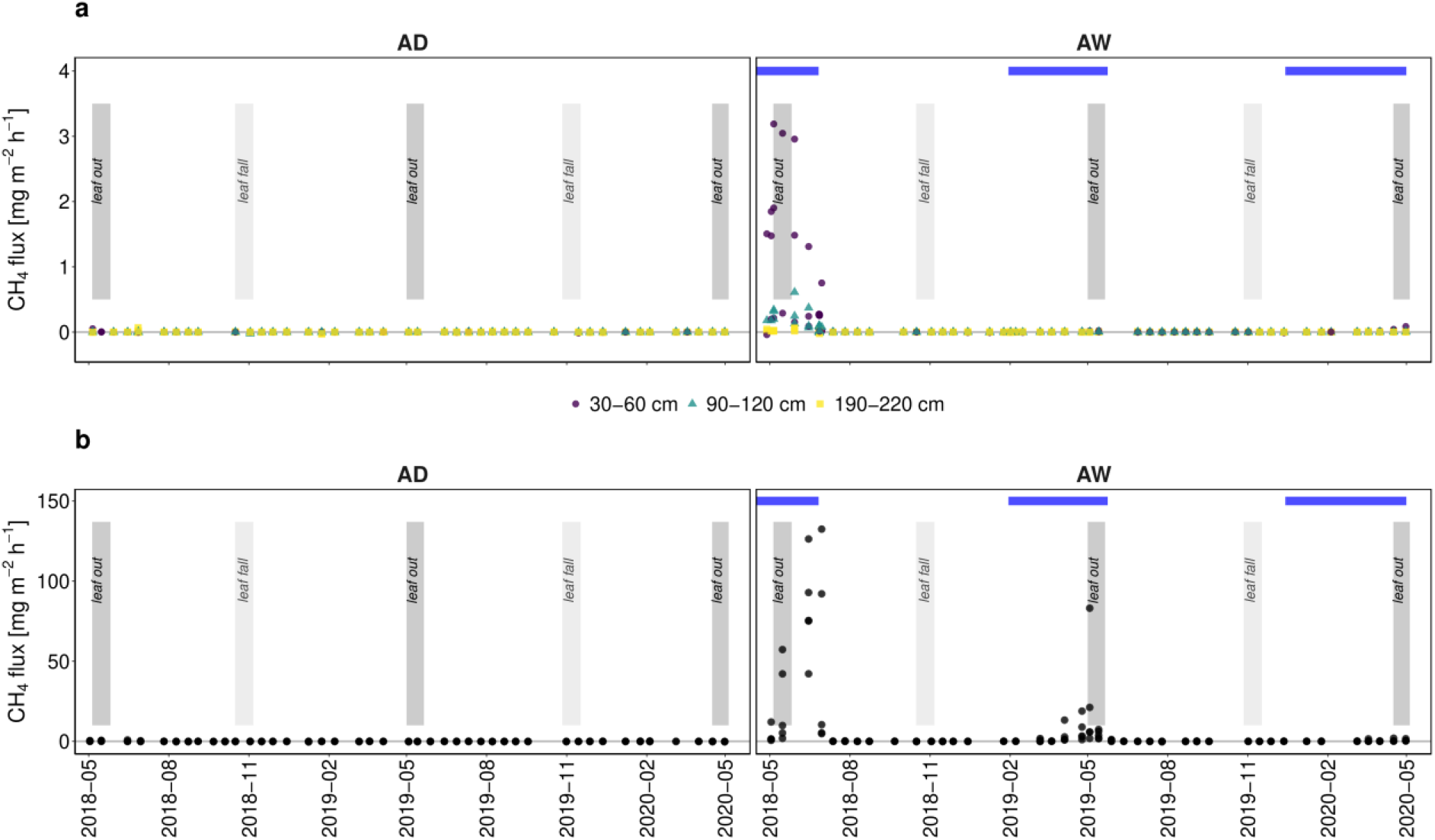
Seasonal course of CH_4_ emissions from **a)** tree stem chambers by their height on the stem and **b)** soil chambers from AD and AW. Points indicate single measurements. Vertical grey bars depict period of leaf out and leaf fall. Horizontal blue bars indicate period of inundation at AW. Note different y-axes on **a** and **b**.

**Fig. 4:**
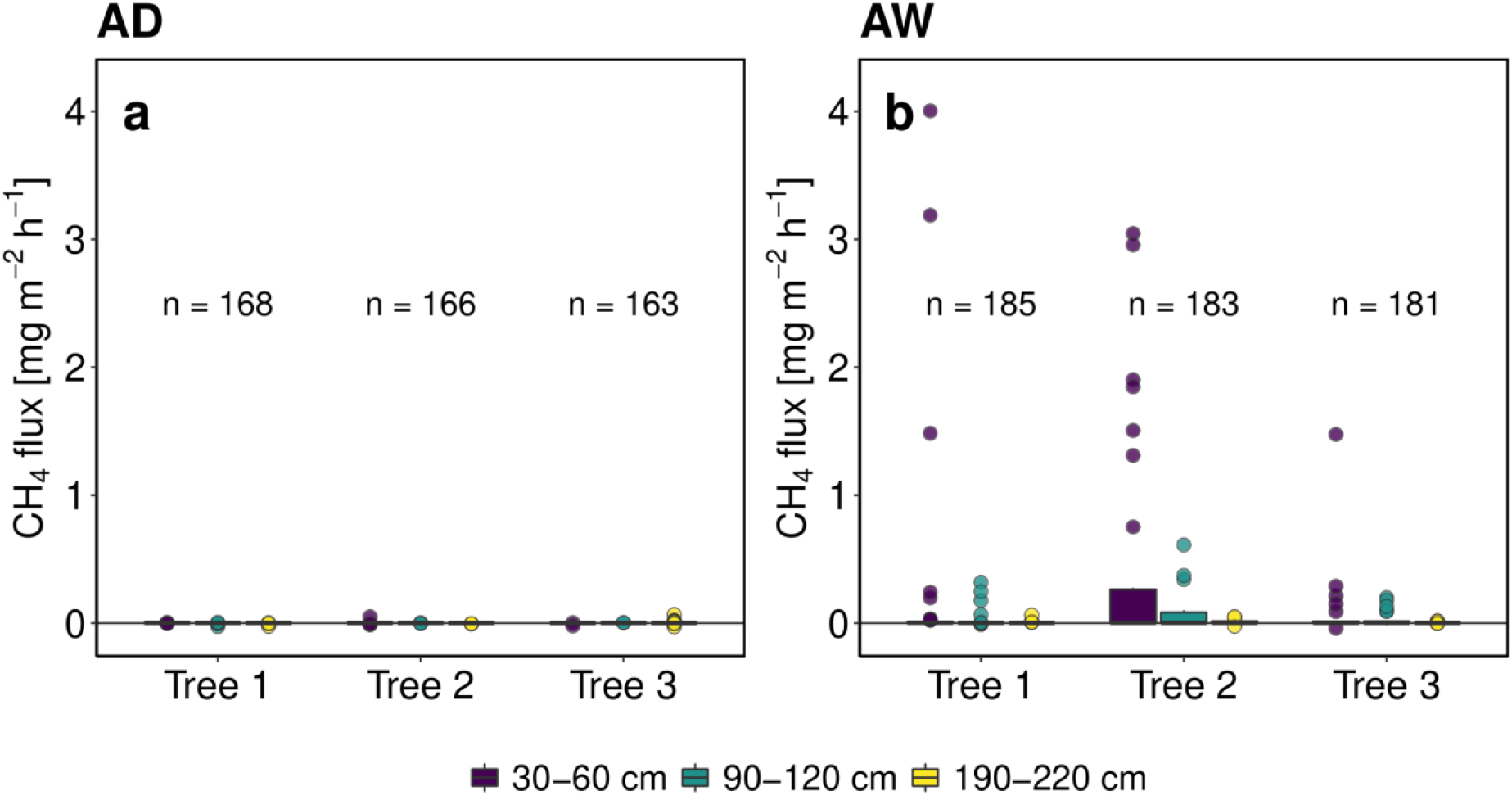
Stem CH_4_ emissions by site **a)** AD and **b)** AW and height of chamber on the stem.

At AW the diameter at breast height (DBH) had a significant impact on stem CH_4_ fluxes (p = 3.2 * 10^−3^). During times when stem CH_4_ emissions could be detected, a weak exponential linear relationship between the height above the stem base (HASB) and stem emissions could be observed (R^2^ 0.12, p = 1.3*10^−6^). Due to the erratic patterns of the stem as well as of the soil CH_4_ emissions, relationships between soil temperature or air temperature were not apparent. However, a significant exponential linear relationship between water level and stem as well as soil CH_4_ emissions was observed at AW (R^2^ = 0.24, p=2.2*10^−16^ and R^2^ = 0.22, p = 1.8 * 10^−11^, respectively). Soil CH_4_ emissions tended to be highest when the water level was just above the soil surface, while stem CH_4_ emissions were higher when the soil was inundated by approximately 25 cm (Fig. 3).

### CH_4_ budgets

Annual CH_4_ budgets differed strongly between stem and soil surfaces and between measuring years at AW. During year one the soil CH_4_ budget for AW was higher than for the second year by a factor of 20 (Table 1).

**Table 1:** Absolute annual balances of stem CH_4_ emissions (±SD) [kg ha^−1^ a^−1^] and the relative share [%] of AD and AW and year one and two. Based on upscaling approaches A1, A2 and A3.

| AD |  |  |  |  |  |  |
| --- | --- | --- | --- | --- | --- | --- |
|  | Year 1 |  |  | Year 2 |  |  |
| approach | soil<sub>absolute</sub> | stem<sub>absolute</sub> | share<sub>stem</sub> [%] | soil<sub>absolute</sub> | stem<sub>absolute</sub> | share<sub>stem</sub> [%] |
| A1 | -2.9±0.7 | 0.002±0.002 | 0.07 | -6.8±0.3 | 0.004±0.001 | 0.06 |
| A2 | -2.9±0.7 | 0.001±0.002 | 0.03 | -6.8±0.3 | 0.004±0.001 | 0.07 |
| A3 | -4.1±5.7 | 0.007±0.063 | 0.17 | -7.8±13.2 | 0.005±0.020 | 0.05 |
| AW |  |  |  |  |  |  |
|  | Year 1 |  |  | Year 2 |  |  |
|  | soil<sub>absolute</sub> | stem<sub>absolute</sub> | share<sub>stem</sub> [%] | soil<sub>absolute</sub> | stem<sub>absolute</sub> | share<sub>stem</sub> [%] |
| A1 | 753±93 | 1.2±0.1 | 0.16 | 51±10 | 0.01±0.001 | 0.02 |
| A2 | 753±93 | 2±0.2 | 0.26 | 51±10 | 0.07±0.002 | 0.14 |
| A3 | 120±358 | 0.17±0.75 | 0.14 | 20±120 | 0.004±0.014 | 0.02 |

All approaches yielded similar results in absolute balances as well as in the shares stem emissions constitute. The overall shares of stem CH_4_ emissions were very low, not exceeding 0.26% in any of the upscaling approaches. Overall, the share of stem-mediated CH_4_ emissions decreased strongly in the second study year, despite decreasing soil CH_4_ emissions. Additionally, shares of stem-mediated CH_4_ emissions at AD are lower than in AW, regardless of the lower soil CH_4_ emissions.

### Leaf fall

The incubation experiment revealed a clear enhancing effect of the addition of alder leaves on the production potential of CH_4_. For incubations of *Soil+Litter_combination_*, CH_4_ production increased by 924% and 774% in AW and AD, respectively, compared to *S_only_+Leaves_only_* (Fig. 5). An increase in CH_4_ production started following a four-day lag phase and peaked after 14 days. Thereafter, the production of CH_4_ decreased gradually and ceased after 30 days of incubation. The production potential in *Soil+Litter_combination_* did not differ significantly between the sites.

**Fig. 5:**
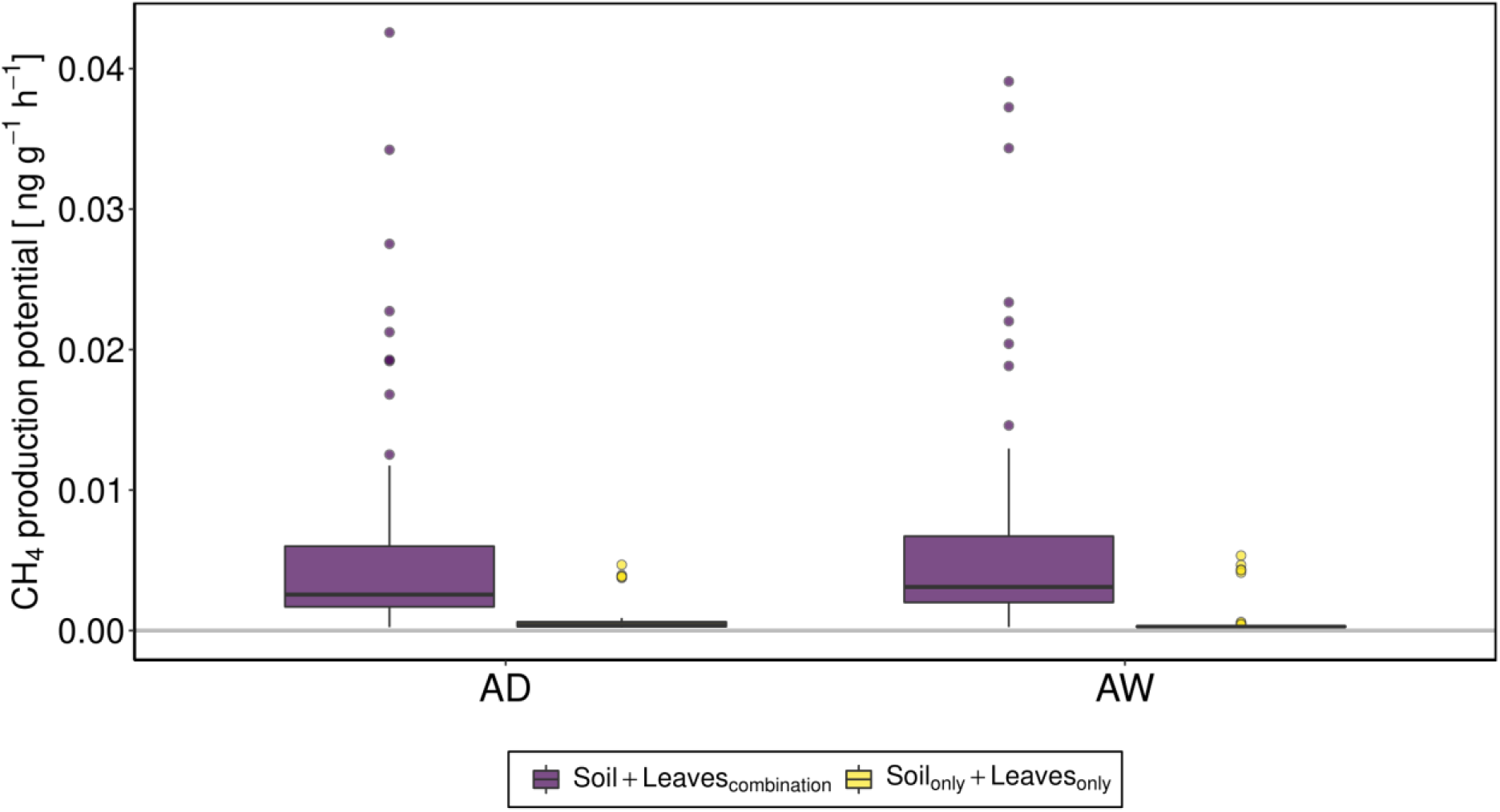
CH_4_ production potential from both incubation treatments. Soil+Leaves_combination_ and Soil_only_ + Leaves_only_ from both sites (AD and AW).

However, soil CH_4_ fluxes at both AD and AW showed no clear increase after leaf fall (Fig. 3). In 2018, CH_4_ fluxes at AW were even slightly lower within four weeks after leaf fall compared to the four weeks prior leaf fall (p = 3.3 * 10^−3^). No significant difference of CH_4_ fluxes before and after leaf fall could be observed at either of the other leaf fall events (AW in 2019, AD in 2018 or in 2019).

## Discussion

### The importance of stem-mediated CH_4_ emissions

Our results show that stems of mature black alder trees act as conduits for CH_4_ and can, at least temporarily, contribute a relevant share of overall ecosystem CH_4_ emissions. Overall, all upscaling approaches gave similarly low shares of stem emissions compared to other studies (Pangala *et al*., 2013, 2015). Only in AD where soil CH_4_ exchange mainly consisted of CH_4_ uptake, the results of the three approaches differed up to one order of magnitude. Even during the period of peak emissions in AW in spring 2018, ecosystem shares of stem CH_4_ emissions were lower than 0.33 % due to the simultaneously very high soil CH_4_ emissions. Interestingly, 87 % of all stem CH_4_ emissions in year one occurred in a two-month period between May 1^st^ and July 1^st^ of 2018 whereas only 7 % of the total soil CH_4_ emissions were detected in the same period (calculated using approach A1). This suggests a potentially transient nature of stem CH_4_ emissions. For future studies it would be interesting to estimate the share of stem-mediated CH_4_ emissions in a constantly inundated alder forest where stem emissions may persist while soil emissions may be hampered by oxidation in the water column (Bastviken *et al*., 2002, 2008). In such environments, the importance of stem-mediated CH_4_ emissions could be much higher. As large parts of inundated swamps are covered by black alder and other wetland tree species, these ecosystems may still be of high importance for the regional CH_4_ budget.

Individual soil CH_4_ fluxes during spring 2018 were extremely high compared to other studies from temperate wetlands (Turetsky *et al*., 2014; Pangala *et al*., 2015). However, on average soil CH_4_ fluxes at AW in this study (4.82 mg m^−2^ h^−1^) fit very well to the approximate mean (4.54 mg m^−2^ h^−1^) for emissions from temperate peatlands (Turetsky et al., 2014).

### Drained vs. undrained alder forest

Despite drought conditions in both study sites, the wet alder forest (AW) on average had distinctively higher CH_4_ emissions from soil and stems than the drained forest. The results therefore support the high importance of water availability or water level for stem CH_4_ emissions (Pitz *et al*., 2018; Barba *et al*., 2019; Schindler *et al*., 2020). Interestingly, stems at AD did not emit CH_4_ and therefore did not offset the sink function of the soil, contrary to what was found at other dry forests (Wang *et al*., 2016; Pitz & Megonigal, 2017).It is possibly that the trees were not connected to the groundwater as rooting depth of black alder varies considerably (Pietzarka & Roloff, 2000), or that no CH_4_ was produced in the soil, due to low carbon content in lower soil layers.

The influence of inundation, drought, leaf out and leaf fall on stem CH_4_ emissions Our data indicates that most likely a combination of inundation, increasing temperatures and increasing physiological activity of vegetation and/or microorganisms leads to higher emission rates of CH_4_ from both soil and stem surfaces. This supports the most important influencing factors for CH_4_ emissions from stems as found in other studies (Barba *et al*., 2018).

In spring 2019 no emissions could be measured at the stem surfaces and soil fluxes showed only a relatively small emission peak that wasof a lower magnitude than the emission maximum in early 2018 (Fig. 3). This may indicate that the drier conditions and shorter period of inundation in spring 2019 compared to 2018 lead to lower CH_4_ production in the soil, and that possibly thresholds regarding the duration of inundation must be exceeded before stem CH_4_ emissions occur. Longer periods of inundation could be related to the time necessary for the depletion of oxygen in the soil. Also, increases in methanogen activity may start with a lag time, as shown in incubation experiments (Sun *et al*., 2012; Ye *et al*., 2015). During low concentrations and production rates of CH_4_ (in the soil) the dominating mode of transport inside the tree is likely to be passive diffusion (Kutschera *et al*., 2016). Hence, it could be assumed that in 2019 when fluxes and presumably soil CH_4_ concentrations were generally lower, the CH_4_ first degassed via the soil-atmosphere interface, because the mode of transport was relatively slow. Contrastingly, in 2018 the CH_4_ concentrations were higher, potentially triggering another mode of transport (Kutschera *et al*., 2016) from the roots to the trunk bypassing potential oxidative layers leading to higher stem fluxes. This, however is based upon the assumption that all CH_4_ is produced in the soil. The uncertainty regarding the transport processes again calls for future detailed parallel investigations of CH_4_ related processes in the soil, pore water and stem. Ideally, also matter turn-over and microbial community should be monitored in direct connection with flux measurements.

Climatic conditions during the study were very variable. generally, drought entirely stopped CH_4_ emissions from both stems and soil but fostered CH_4_ uptake in both sites. Conditions at AW even changed so drastically that despite previously showing strong emissions in June 2018 the soil repeatedly turned to a sink in the following summer with a maximum uptake of −0.3 mg m^−2^ h^−1^ in July 2019, similar to the uptake rate at AD. This effect could be explained by a high abundance of methanotrophic Bacteria and Archaea at AW (Weil *et al*., 2020). Since global warming increases climate variability and the likelihood for extreme events like drought and strong precipitation (King & Karoly, 2017), the importance of stem CH_4_ emissions could be changed drastically in the future. Thus, long term monitoring and understanding of the driving factors in variable environmental conditions is still important to estimate the impact of stem-mediated emissions.

The observed emission peaks took place parallel to leaf out of the black alder, indicating that through a rise in activity and transpiration rates inside the tree possibly more CH_4_ could enter the tree. Sap flow as a meter for transpiration can act as an explanatory variable for stem CH_4_ emissions (Barba *et al*., 2019). Further, the results of Barba et al. (2019) suggest that increasing temperatures also directly lead to an increase in stem CH_4_ emissions. Thus, it is likely that a combination of sufficient water availability, sap flow intensity and rising temperatures led to peak stem CH_4_ emission in our study. Aside from sap flow intensity, stem CO2 emissions could be used as an indicator for respiration and hence indirectly for physiological activity of the tree (Hölttä & Kolari, 2009; Machacova *et al*., 2019), that may explain stem CH_4_ emissions (Barba *et al*., 2017). However, in our study stem CO2 emissions at AW could only explain 1% of all CH_4_ fluxes larger than 0.1 mg m^−2^ h^−1^ (tested with linear models, data not shown). Thus, stem CH_4_ emissions in our study seem to be independent from physiological activity of the tree. This is further supported by our finding that fluxes did not vary over the course of the day (data not shown).

Additionally, our data suggests that individual differences in stem CH_4_ emissions between trees can be large, despite all trees being of similar size and of the same species (Fig. 4). Thus, other factors that were not considered in this study, both concerning the soil and the tree, may be of importance for stem CH_4_ emissions. Microbiological community and microtopography may influence local methanogenesis and, thus, pore water CH_4_ concentration in the rooting zone of the tree (Pangala *et al*., 2015; Terazawa *et al*., 2015). Physiological activity inside the tree (Zeikus & Ward, 1974; Covey *et al*., 2012) or morphological parameters such as wood density could be responsible for significant differences between individual trees (Pangala *et al*., 2013). Here it would be useful to expand the study over the entire stand and sample for stem diameter and wood specific density.

Finally, as indicated by our incubation experiment leaf fall holds the potential to increase CH_4_ production, probably by providing fresh substrate for methanogenesis. However, the results from the incubation are not directly transferable to the field, as fluxes measured in the field did not increase in the weeks after leaf fall. In the field, both temperature and water availability may have been limiting factors for methanogenesis despite the input of potential substrate. Hence, it is unlikely that in temperate regions leaf fall alone leads to CH_4_ emission peaks. Therefore, the long-term effects of leaves on CH_4_ emissions should be studied in the future through combination of multiple-site leaf-exclusion and incubation/mesocosm experiments. Especially alder stands that are directly connected with constant water bodies such as rivers and lakes – as is often the case in north and central Europe (Claessens *et al*., 2010) – hold the potential of being a constant overlooked mediator of CH_4_.

### Conclusion

Data from this and other studies show that forests must not be overlooked in global CH_4_ budgets. However, the importance of stem-mediated CH_4_ emissions may vary considerably between individual ecosystems, even if dominated by the same tree species. The present study shows that the variability of stem and soil CH_4_ emissions cannot be explained by a single influential factor and likely differs strongly across ecosystems. Further, the interplay of all variables involved demands future investigations. Although our findings suggest the soil to be the origin of the CH_4_ coming from the stems the observed variability of this emission path and the unknown transport mechanism inside the tree clearly ask for more mechanistic research within this topic.

## Acknowledgements

The authors thank field technicians, student assistants and volunteers for supporting the measurement campaigns and maintaining the equipment.

## Author Contributions

Design of the research: D.K., A.G and G.J.; field work was conducted by: D.K., I.S.; data analysis, collection, or interpretation: D.K., A.G., I.S., G.J.; writing the manuscript: D.K., A.G., G.J.

## Data availability

Data is accessible via PANGAEA link: https://issues.pangaea.de/browse/PDI-25060

## References

1. Barba J, Bradford MA, Brewer PE, Bruhn D, Covey K, van Haren J, Megonigal JP, Mikkelsen TN, Pangala SR, Pihlatie M, et al. 2018. Methane emissions from tree stems: a new frontier in the global carbon cycle. New Phytologist 222: 18–28.

2. Barba J, Poyatos R, Vargas R. 2017. Automated CO_2_,CH_4_ and N_2_O fluxes from tree stems and soils: magnitudes, temporal patterns and drivers. In: Fall Meeting, AGU, New Orleans, LA, USA, 11–15 December.

3. Barba J, Poyatos R, Vargas R. 2019. Automated measurements of greenhouse gases fluxes from tree stems and soils: magnitudes, patterns and drivers. Scientific Reports 9: 1–13.

4. Bartlett KB, Harriss RC, Andersen BL, B G, Leip A, Rembges D, Bergamaschi P, Lubina C, Fischer H, Fisher RE, et al. 2006. Methane emissions from terrestrial plants under aerobic conditions. Nature 13: 445–461.

5. Bastviken D, Cole JJ, Pace ML, Van de-Bogert MC. 2008. Fates of methane from different lake habitats: Connecting whole-lake budgets and CH_4_ emissions. Journal of Geophysical Research: Biogeosciences 113: 1–13.

6. Bastviken D, Ejlertsson J, Tranvik L. 2002. Measurement of methane oxidation in lakes: A comparison of methods. Environmental Science and Technology 36: 3354–3361.

7. Bižić M, Klintzsch T, Ionescu D, Hindiyeh MY, Günthel M, Muro-Pastor AM, Eckert W, Urich T, Keppler F, Grossart HP. 2020. Aquatic and terrestrial cyanobacteria produce methane. Science Advances 6: 1–10.

8. Claessens H, Oosterbaan A, Savill P, Rondeux J. 2010. A review of the characteristics of black alder (*Alnus glutinosa* (L.) Gaertn.) and their implications for silvicultural practices. Forestry 83: 163–175.

9. Covey KR, Wood SA, Warren RJ, Lee X, Bradford MA. 2012. Elevated methane concentrations in trees of an upland forest. Geophysical Research Letters 39: 1–6.

10. Gauci V, Gowing DJG, Hornibrook ERC, Davis JM, Dise NB. 2010. Woody stem methane emission in mature wetland alder trees. Atmospheric Environment 44: 2157–2160.

11. Günther A, Jurasinski G, Albrecht K, Gaudig G, Krebs M, Glatzel S. 2017. Greenhouse gas balance of an establishing Sphagnum culture on a former bog grassland in Germany. Mires and Peat 20: Article 02.

12. Günther A, Jurasinski G, Huth V, Glatzel S. 2014. Opaque closed chambers underestimate methane fluxes of *Phragmites australis* (Cav.) Trin. ex Steud. Environmental Monitoring and Assessment 186: 2151–2158.

13. Hölttä T, Kolari P. 2009. Interpretation of stem CO2 efflux measurements. Tree Physiology 29: 1447–1456.

14. Joabsson A, Christensen TR. 2001. Methane emissions from wetlands and their relationship with vascular plants: an Arctic example. Global Change Biology 7: 919–932.

15. Jurasinski G, Ahmad S, Anadon-Rosell A, Berendt J, Beyer F, Bill R, Blume-Werry G, Couwenberg J, Günther A, Joosten H, et al. 2020. From Understanding to Sustainable Use of Peatlands: The WETSCAPES Approach. Soil Systems 4: 1–27.

16. Jurasinski G, Koebsch F, Günther A, Beetz S. 2014. R package Flux.

17. King AD, Karoly DJ. 2017. Climate extremes in Europe at 1.5 and 2 degrees of global warming. Environmental Research Letters 12: 1–9.

18. Kutschera E, Khalil A, Rice A, Rosenstiel T. 2016. Mechanisms of methane transport through *Populus trichocarpa*. Biogeosciences Discussions 60: 1–17.

19. Livingston GP, Hutchinson GL. 1995. Enclosure-based measurement of trace gas exchange: applications and sources of error. In: Matson PA, Harris RC, eds. Biogenic Trace Gases: Measuring Emissions from Soil and Water. 14–53.

20. Machacova K, Bäck J, Vanhatalo A, Halmeenmäki E, Kolari P, Mammarella I, Pumpanen J, Acosta M, Urban O, Pihlatie M. 2016. *Pinus sylvestris* as a missing source of nitrous oxide and methane in boreal forest. Scientific Reports 6: 2–10.

21. Machacova K, Vainio E, Urban O, Pihlatie M. 2019. Seasonal dynamics of stem N_2_O exchange follow the physiological activity of boreal trees. Nature Communications: 1–13.

22. Maier M, Machacova K, Lang F, Svobodova K, Urban O. 2017. Combining soil and tree-stem flux measurements and soil gas profiles to understand CH 4 pathways in *Fagus sylvatica* forests. Journal of Plant Nutrition and Soil Science.

23. Melton JR, Wania R, Hodson EL, Poulter B, Ringeval B, Spahni R, Bohn T, Avis CA, Beerling DJ, Chen G, et al. 2013. Present state of global wetland extent and wetland methane modelling: Conclusions from a model inter-comparison project (WETCHIMP). Biogeosciences 10: 753–788.

24. Pangala SR, Enrich-Prast A, Basso LS, Peixoto RB, Bastviken D, Hornibrook ERC, Gatti L V., Marotta H, Calazans LSB, Sakuragui CM, et al. 2017. Large emissions from floodplain trees close the Amazon methane budget. Nature 552: 230–234.

25. Pangala SR, Gowing DJ, Hornibrook ERC, Gauci V. 2014. Controls on methane emissions from *Alnus glutinosa* saplings. New Phytologist 201: 887–896.

26. Pangala SR, Hornibrook ERC, Gowing DJ, Gauci V. 2015. The contribution of trees to ecosystem methane emissions in a temperate forested wetland. Global Change Biology 21: 2642–2654.

27. Pangala SR, Moore S, Hornibrook ERC, Gauci V. 2013. Trees are major conduits for methane egress from tropical forested wetlands. New Phytologist 197: 524–531.

28. Pietzarka U, Roloff A. 2000. Alnus glutinosa. In: Enzyklopädie der Holzgewächse. 1–16.

29. Pitz S, Megonigal JP. 2017. Temperate forest methane sink diminished by tree emissions. New Phytologist 214: 1432–1439.

30. Pitz SL, Megonigal JP, Chang CH, Szlavecz K. 2018. Methane fluxes from tree stems and soils along a habitat gradient. Biogeochemistry 137: 307–320.

31. R development core team. 2020. R 3.6.4.

32. Rusch H, Rennenberg H. 1998. Black alder (*Alnus glutinosa* (L.) Gaertn.) trees mediate methane and nitrous oxide emission from the soil to the atmosphere. Plant and Soil 201: 1–7.

33. Saunois M, Bousquet P, Poulter B, Peregon A, Ciais P, Canadell JG, Dlugokencky EJ, Etiope G, Bastviken D. 2016. The global methane budget 2000 – 2012. Earth System Scientific Data 8: 697–751.

34. Schäfer A, Barthelmes A, Joosten H, Kaffke A, Koska I, Schröder J, Succow M. 2005. Erlenaufforstung auf wiedervernässten Niedermooren (ALNUS-Leitfaden).

35. Schindler T, Mander Ü, Machacova K, Espenberg M, Krasnov D. 2020. Short-term flooding increases CH_4_ and N_2_O emissions from trees in a riparian forest soil-stem continuum. Scientific Reports 10: 1–10.

36. Schröder P. 1989. Characterization of a thermo-osmotic gas transport mechanism in *Alnus glutinosa* (L.) Gaertn. Trees 3: 38–44.

37. Siegel AF. 1982. Robust regression using repeated medians. Biometrika 69: 242–244.

38. Sun CL, Brauer SL, Cadillo-Quiroz H, Zinder SH, Yavitt JB. 2012. Seasonal changes in methanogenesis and methanogenic community in three peatlands, Newyork State. Frontiers in Microbiology 3: 1–8.

39. Terazawa K, Ishizuka S, Sakata T, Yamada K, Takahashi M. 2007. Methane emissions from stems of *Fraxinus mandshurica* var. japonica trees in a floodplain forest. Soil Biology and Biochemistry 39: 2689–2692.

40. Terazawa K, Yamada K, Ohno Y, Sakata T, Ishizuka S. 2015. Spatial and temporal variability in methane emissions from tree stems of *Fraxinus mandshurica* in a cool-temperate floodplain forest. Biogeochemistry 123: 349–362.

41. Turetsky MR, Kotowska A, Bubier J, Dise NB, Crill P, Hornibrook ERC, Minkkinen K, Moore TR, Myers-Smith IH, Nykänen H, et al. 2014. A synthesis of methane emissions from 71 northern, temperate, and subtropical wetlands. Global Change Biology 20: 2183–2197.

42. Wang ZP, Gu Q, Deng FD, Huang JH, Megonigal JP, Yu Q, Lü XT, Li LH, Chang S, Zhang YH, et al. 2016. Methane emissions from the trunks of living trees on upland soils. New Phytologist 211: 429–439.

43. Weil M, Wang H, Bengtsson M, Köhn D, Günther A, Jurasinski G, Couwenberg J, Negassa W, Zak D, Urich T. 2020. Long-term rewetting of three formerly drained peatlands drives congruent compositional changes in pro- and eukaryotic soil microbiomes through environmental filtering. Microorganisms 8.

44. Ye R, Doane TA, Morris J, Horwath WR. 2015. The effect of rice straw on the priming of soil organic matter and methane production in peat soils. Soil Biology and Biochemistry 81: 98–107.

45. Zeikus JG, Ward J.C. 1974. Methane Formation in Living Trees - A Microbial Origin. Science 184: 1181–1183.

